# Culture-enriched metagenomics enables genome-resolved detection of low abundance ESKAPE and *Vibrio* pathogens in coastal habitats

**DOI:** 10.64898/2026.05.14.725077

**Authors:** Jia Yee Ho, Dalong Hu, Deborah Yebon Kang, Clarence Bo Wen Sim, Winona Wijaya, Yann Felix Boucher

**Author notes:** **Corresponding author:** Yann Felix Boucher. Equal contribution and first authorship (Jia Yee Ho, Dalong Hu).

## Abstract

Coastal marine environments are increasingly recognised as reservoirs of antimicrobial-resistant (AMR) pathogens. However, it remains challenging to recover high-quality genomes of clinically relevant bacteria present at low abundance from complex natural systems. Here, we applied culture-enriched metagenomics to systematically track the diversity and dynamics of major AMR pathogens within the coastal marine system of St. John’s Island, Singapore, as a model ecosystem for pathogen surveillance. Selective media-based enrichment recovered 773 metagenome-assembled genomes (MAGs) from 92 multi-matrix environmental samples, which includes coastal water, sediment, and seaweed, capturing diverse AMR ESKAPE and *Vibrio* species. Distinct bacterial signatures and dispersal patterns were observed in each niche, for example, microbes that signal human impact was detected at the beach, while fish-associated pathogens were present at the aquaculture facility outlet. Notably, the high-quality MAGs enabled subspecies-level identification and supported the AMR gene detection across six distinct coastal habitats. Detailed differences in the recovery of specific pathogens across enrichment media were also identified, demonstrating the method’s efficacy in finding media suitable for surveillance of specific organisms, such as deciding between liquid or solid formulations. MAGs recovered from culture-enriched metagenomics were highly similar to genomes obtained from pure isolates, as demonstrated for *Klebsiella pneumoniae*. The preserved culture-enriched stocks were capable of recovering organisms of interest when individual isolates were required for further study. Overall, our findings highlight the utility of culture-enriched metagenomics as a cost-effective, sensitive approach to uncovering the genomic landscape of pathogens with environmental reservoirs, with implications for AMR surveillance and ecological risk assessment.

## Introduction

Antimicrobial resistance (AMR) poses a major threat to public health that could result in morbidity, mortality and significant economic losses. In 2019, it was estimated that over 1.2 million deaths were associated with AMR infections across 204 countries (Murray et al., 2022). It was also projected that the deaths caused by AMR will overtake cancer and other non-communicable diseases, including diabetes, to reach 10 million in 2050 (Jim O’Neill, 2014). In the WHO Bacterial Priority Pathogens List 2024 (BPPL-2024), carbapenem resistant *Acinetobacter baumannii* (CRAB), carbapenem-resistant Enterobacterales (CRE) and third generation cephalosporin-resistant Enterobacterales (3GCRE) are classified in the critical priority category (World Health Organization, 2024). Meanwhile, vancomycin-resistant *Enterococcus faecium* and carbapenem-resistant *Pseudomonas aeruginosa* (CRPA) are listed in the high priority category (World Health Organization, 2024). The acronym “ESKAPE” was introduced by the Infectious Diseases Society of America to refer to a group of pathogens that have a high capacity to develop AMR, namely *<u>E</u>nterococcus faecium*, *<u>S</u>taphylococcus aureus*, *<u>K</u>lebsiella pneumoniae*, *<u>A</u>cinetobacter baumannii*, *<u>P</u>seudomonas aeruginosa*, and *<u>E</u>nterobacter* species (Rice, 2008). These pathogens pose a formidable global challenge due to their widespread prevalence and resistance, spreading silently within the environment, communities and healthcare settings. Because infections caused by resistant strains often mirror those from susceptible strains and detection is challenging. Without robust surveillance systems, the detection of emerging resistance trends is frequently delayed, leading to postponed infection control measures, inappropriate antimicrobial use, and larger outbreaks (Ventola, 2015, Laxminarayan et al., 2013). Studies in Singapore and elsewhere have shown that surface waters, especially near agricultural or hospital-influenced areas, harbour AMR bacteria and genes, posing human exposure risks and highlighting potential transmission between humans and the environment (Goh et al., 2023, Goh et al., 2024, Leonard et al., 2015, Leonard et al., 2018a, Leonard et al., 2018b, O’Flaherty et al., 2019, Schijven et al., 2015, Yuan et al., 2024, Ho et al., 2021).

AMR bacteria surveillance is traditionally based on culturing and quantitative polymerase chain reaction (qPCR) approaches. However, antimicrobial resistance genes (ARGs) detected through qPCR do not contain the information associating them to host bacteria strains. To gain such insight, a culturing approach followed by whole genome sequencing (WGS) is usually used, thus informing both the AMR phenotype and the host genotype (Duggett et al., 2020). To bypass the intensive process of isolating pure bacterial colonies, metagenomic sequencing enables comprehensive profiling of entire microbial communities directly from environmental matrices. It serves as a robust tool to discover phylogenetically novel marine bacteria (Ho et al., 2025b). However, a significant limitation of current culture-independent metagenomic surveillance is the high concentration of non-target ‘background’ microbial DNA (Schrader et al., 2012). This often results in insufficient sequencing depth for low-abundance pathogens or fastidious organisms amidst this environmental ‘noise’ (Zaheer et al., 2018). Consequently, clinically significant resistant variants often go undetected or are underestimated, leading to an incomplete understanding of the environmental resistome, such as mobile genetic elements, and missed opportunities for early intervention in emerging outbreaks (Partridge et al., 2018). A previous study demonstrated that while taxonomic profiles stabilise at relatively low sequencing depths, recovering the full richness of AMR gene families requires at least 12 Gbp per sample (Gweon et al., 2019). Furthermore, even at a depth of 60 Gbp, the complete allelic diversity of AMR genes remains elusive in complex matrices like river sediment and pig caeca, where target pathogens often exist in low abundance (Gweon et al., 2019).

Furthermore, the resolution power of WGS for epidemiological surveillance can only be approached by metagenomics if high-quality metagenome assembled genomes (MAGs) can be obtained from metagenomic data. Some metagenomics studies have detected ARGs in MAGs within environmental samples, but so far only in very small numbers and from locations with high exposure to a dense human population such as transits systems, wastewater, and water purification facilities (Stamps and Spear, 2020, Magnúsdóttir et al., 2023, Abdulkadir et al., 2024). Such inefficient and costly testing is challenging to implement in epidemiological surveillance, which demands large-scale, routine operations.

Culture-enriched metagenomics is an adaptation of metagenomics and WGS designed to achieve greater genomic recovery of species of interest, which has so far been used primarily in food sample pathogen detection (Pronyk et al., 2023, Hyeon et al., 2018, Kocurek et al., 2023, Ottesen et al., 2020, Townsend et al., 2020). Recently, applications have expanded to human, environmental and plant-associated microbial communities, including AMR surveillance, which outperforms metagenomics in terms of quality of MAGs or detection of ARGs (Raymond et al., 2019, Whelan et al., 2020, Ottesen et al., 2022, Fu et al., 2023, Li et al., 2023, Zhang et al., 2022, Yu et al., 2024). While some of these previous studies successfully recovered MAGs, they often remained limited to single-environmental matrix, focusing exclusively on either water, soil, desert soil, as well as wastewater and its receiving river, which precludes a comparative understanding of lineages distribution across interconnected niches like water, sediment, and macroalgae. Additionally, there remains a lack of direct genomic validation comparing the computationally derived MAGs to pure isolates obtained from the same enrichment stocks, which is critical for confirming the accuracy of strain-level surveillance.

St. John’s Island (SJI), formerly known as Pulau Sekijang, spans 39 hectares and is located approximately 6.5 km south of mainland Singapore. The island has a rich history, in the past housing a British-run quarantine centre for cholera, smallpox, and the plague for nearly a century; a detention centre for political prisoners; and an opium treatment centre (National Parks Board). SJI is now connected to Lazarus Island, which was renamed when a lazaretto (quarantine hospital) was built there in 1899. Today, SJI with Lazarus Island, forms a complex with two other islands, Pulau Seringat and Pulau Kias. These uninhabited islands serve as high-traffic recreational destinations for both domestic and international visitors. Anthropogenic activities on the island are primarily characterised by water-based recreation (e.g., swimming, kayaking) and shore-based activities such as camping and trekking. The sampling sites include a secluded lagoon within the intertidal zone and an extensive sandy beach on the Lazarus Island, both of which are high-traffic areas for swimming. While these islands are primarily day-trip destinations, limited overnight accommodations are available.

The island was selected as a model site to pilot the environmental surveillance process, as it displays some recreational human activities coupled with well-preserved natural environments. It is uniquely positioned as a diverse ecosystem with minimal direct exposure to antimicrobials. Importantly, all three One Health domains (human, animal, and the environment) are represented in the SJI coastal environment. As microbial pollution does not change monotonically from one location to the other, but is dictated by local waste releases and tidal mixing (Ho et al., 2021), sampling at different locations of the island is required to determine the efficacy of surveillance. Water and sediment samples were collected at six locations on SJI, with seaweed samples obtained at four of these sites.

To date, no studies have utilised culture-enriched metagenomics to simultaneously recover high-quality MAGs across diverse environmental matrices and characterise their associated ARGs. Furthermore, the genomic congruence between computationally assembled MAGs and pure isolates derived from the same enrichment stocks has not been rigorously validated. In this study, we present an integrated approach combining culture-enriched metagenomics with comprehensive bioinformatics analyses to identify AMR bacteria and their associated ARGs in environmental samples from St. John’s and Lazarus Island (**Figure *1***). We demonstrate that this approach enables robust genome recovery with subspecies-level resolution, as well as detection of AMR lineages in complex environmental samples. The method also enables pre-emptive surveillance, a proactive public health strategy designed to identify and assess the risk of potential pathogens before a widespread outbreak occurs, often by monitoring spillover signals in wildlife or environmental reservoirs. We also show that culture-enriched metagenomics can identify bacterial signatures linked to niche selection and anthropogenic impact. Our pipeline recovered 1,124 MAGs in total, of these, 773 MAGs were derived from 92 enrichment samples (800.49 Gbp), compared to 351 MAGs from six direct coastal water metagenomic datasets (11.50 Gbp). For low-abundance pathogens, direct metagenomics showed extremely limited sensitivity for detection in coastal environments. Conversely, our culture-enriched metagenomics recovered high-quality MAGs capturing diverse pathogens, including all ESKAPE and major pathogenic *Vibrio* species. Notably, from culture-enriched metagenomics, 610 MAGs were species-resolved MAGs and of these, 383 MAGs (62.8%) were identified as known pathogens. We observed distinct patterns of pathogen recovery, revealing the taxonomic biases imposed by different enrichment media. We also found that the MAGs recovered from culture-enrichment, specifically *K. pneumoniae*, showed high genomic congruence with pure isolates recovered from the same enrichment stocks. This demonstrates that the integrated approach provides the high-resolution surveillance data necessary for environmental health monitoring.

**Figure 1.**
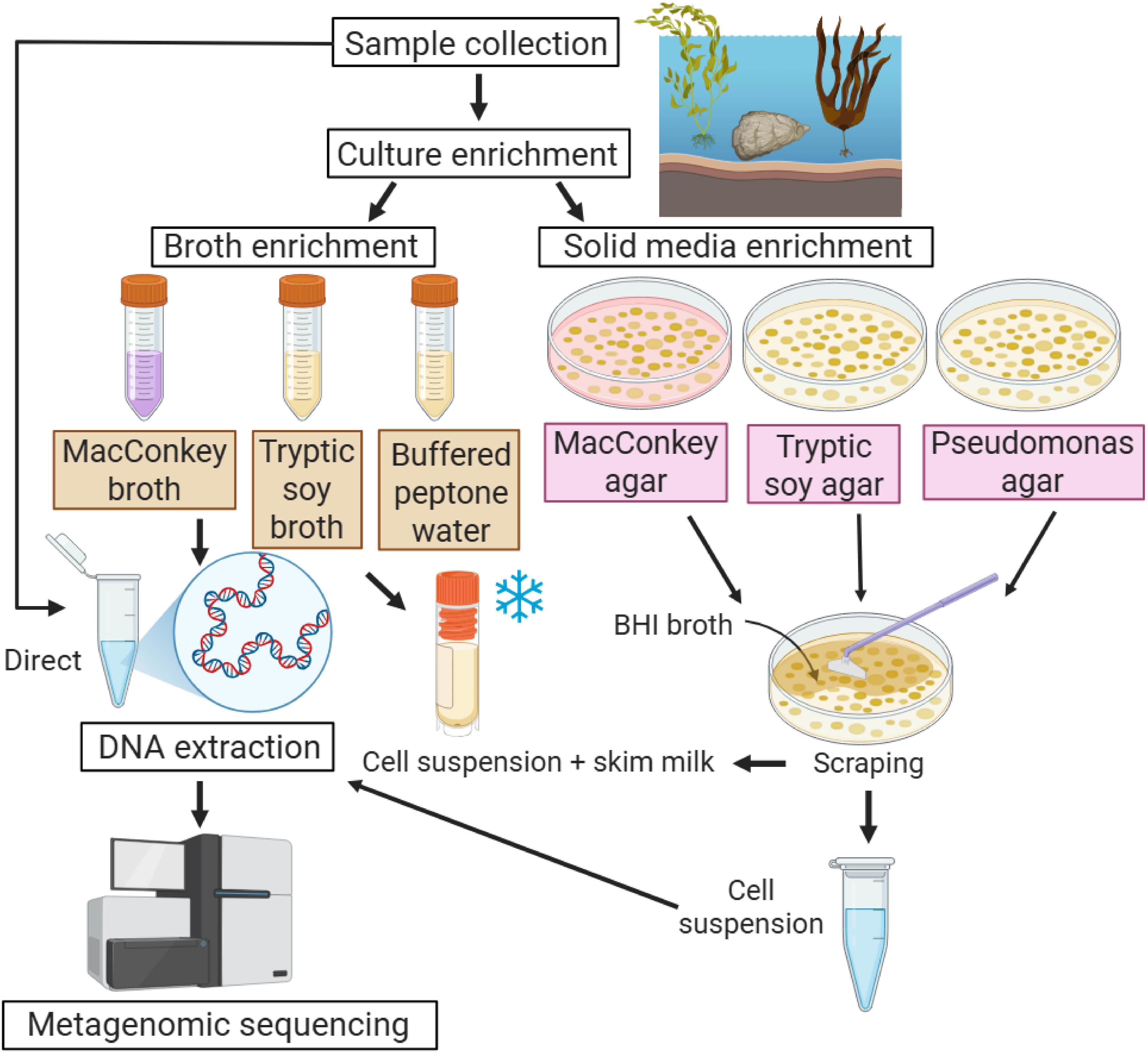
Pipeline of culture-enriched metagenomics for coastal water, macroalgae (seaweed), and sediment samples collected from St John’s Island (SJI). Samples were subjected to either broth or solid media enrichment prior to metagenomic sequencing. Broth and solid media included MacConkey (MacA and MacB) and tryptic soy media (TSA and TSB), in addition to Buffered Peptone Water (BPW) and Pseudomonas agar (PseuA). Solid media enrichment required an additional colony-scraping step using BHI broth to obtain cell suspensions. DNA extraction was performed directly from the original samples (for direct metagenomics) and from the enrichment-derived cell suspensions. Cell suspensions from both solid and liquid enrichments were mixed with skim milk and stored at −80°C for long-term preservation.

## Materials and methods

### Study site and sample collection

St. John’s Island (SJI), Singapore, hosts diverse natural habitats and abundant wildlife, including heritage trees, birds, reptiles, amphibians, insects, fish, crustaceans, molluscs, coral reefs, and seaweed. A small patch of mangroves, dominated by Bakau Pasir (*Rhizophora stylosa*) and Tumu (*Bruguiera gymnorhiza*), forms a living “seawall” for the coast at part of the island (National Parks Board) (**Figure *2***). The western reefs of SJI is part of the Sisters’ Islands Marine Park, Singapore’s first marine protected area. Located within the island is the St. John’s Island National Marine Laboratory (SJINML), Singapore’s only offshore marine research facility under the National Research Infrastructure (NRI) scheme, managed by the National University of Singapore (NUS), to support marine science research that meets strategic national needs (St. John’s Island National Marine Laboratory). SJINML also houses a public gallery at its Marine Park Outreach and Education Centre, with a mangrove mesocosm and operational facility growing corals to complement outreach programmes (National Parks Board).

**Figure 2.**
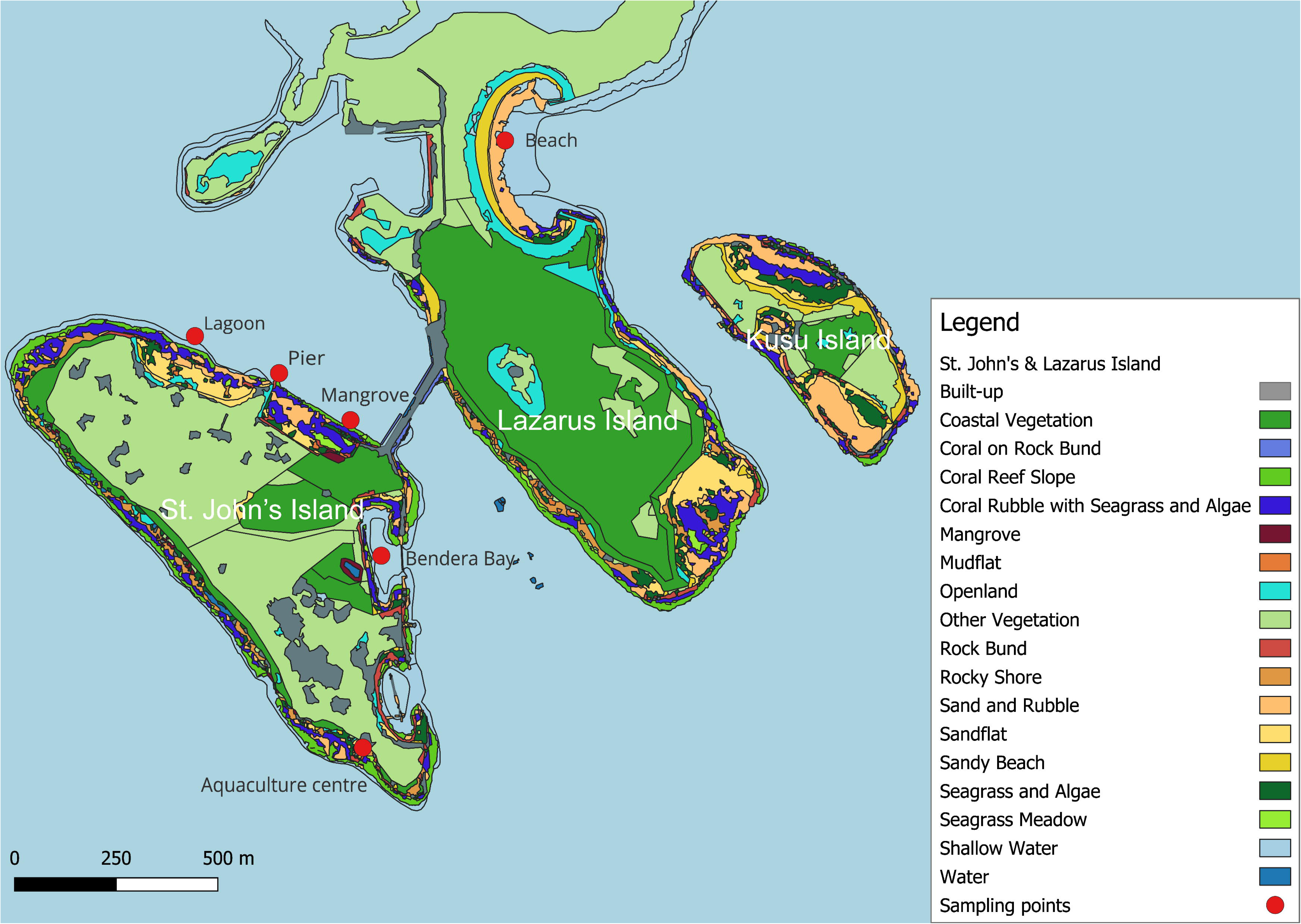
Map of St. John’s and Lazarus Islands with sampling points indicated by red circles. All points include water and sediment samples; seaweed samples were collected only at the ‘Bendera Bay’, ‘Pier’, ‘Lagoon’, and ‘Beach’. The map was created using the Free and Open Source QGIS. The NParks “Parks and Nature Reserves map” and “Coastal and Marine Habitat map” were retrieved from Singapore’s open data portal (National Parks Board, 2023a, National Parks Board, 2023b)

Water and sediment samples were collected from six different locations at SJI (GPS: 1.216595, 103.850151) in August 2022 at low tide. The sampling locations were “Lagoon”, “Pier”, “Mangrove”, “Bendera Bay”, “Aquaculture centre” at SJI and “Beach” at Lazarus Island. At four of the six sampling locations (“Lagoon”, “Pier”, “Bendera Bay”, “Beach”), macroalgae (comprising *Caulerpa* spp., *Gracilaria* spp., *Sargassum* spp., *Ulva* spp., *Padina* spp.) were collected and categorised as the ‘seaweed’ matrix for subsequent analysis. The details in **Figure *2*** was retrieved from Singapore’s open data portal (National Parks Board, 2023a, National Parks Board, 2023b). Sampling was permitted under NParks research permits NP/RP23-060a and NP/RP26-024.

### Bacterial enrichment

The samples were enriched in either broth or agar for cultivation of bacteria. Three different types of agar was used for agar enrichment, namely MacConkey (MacA, Merck Millipore, Darmstadt, Germany), Pseudomonas (PseuA, BD, New Jersey, USA) and tryptic soy agar (TSA, Merck Millipore, Darmstadt, Germany). Sediment samples were subjected to a brief rinsing with sterile MilliQ water. Thereafter, 1 g of the sediment was resuspended in 10 mL of PBS. After vortexing, the suspension was plated directly onto the agar. Seaweed samples were rinsed briefly using sterile MilliQ water and 1 strand of seaweed added to PBS. After brief vortexing, a 1 hour shaking at 180 rpm at room temperature was carried out. The suspension was then plated directly onto the agar. For water samples, a 1 mL and 10 mL was filtered onto 0.22 µm sterile CN membrane filter (Sartorius, Göttingen, Germany) and placed onto PseuA (1 mL), TSA (1 mL) and MacA (10 mL), respectively. All the agar were then incubated overnight at 37°C.

Three enrichment broth media were utilised: MacConkey broth (MacB; Merck Millipore, Darmstadt, Germany), buffered peptone water (BPW; Merck Millipore, Darmstadt, Germany), and tryptic soy broth (TSB; Oxoid, Hampshire, UK). For processing, 1 g of sediment or 1 strand of seaweed were rinsed with sterile MilliQ water and added separately to the respective broths. Simultaneously, 600 mL of water was filtered through a 0.22 µm sterile CN membrane filter (Sartorius, Göttingen, Germany), the resulting membrane was folded in half and submerged in the respective broth. All cultures were then incubated overnight at 37°C with shaking at 220 rpm.

All the bacterial colonies grown on the agar after overnight incubation were then harvested by adding 1.5 mL of brain-heart infusion (BHI, BD, New Jersey, USA) broth and scraping. Aliquots of the cell suspension were then stored for DNA extraction, and 0.5 mL of the aliquot was stored in 20% skim milk for long term storage in −80°C. On the other hand, the cell suspension from the liquid media was aliquoted after overnight incubation and stored for DNA extraction. Similarly, 0.5 mL of the aliquot was stored in 20% skim milk for long-term storage in −80°C.

### DNA extraction and sequencing

For direct metagenomics, DNeasy PowerWater Kit (Qiagen, Venlo, The Netherlands) was used to isolate genomic DNA from the 0.22 µm CN membrane filter (Sartorius, Göttingen, Germany) according to the manufacturer’s protocol. However, seaweed and sediment samples yielded DNA concentrations below the required threshold for library construction, consequently, direct metagenomics was omitted for these matrices. For culture-enriched metagenomics, the cell suspension from agar or broth culture were then subjected to DNA extraction using QIAamp PowerFecal Pro DNA Kit (Qiagen, Venlo, The Netherlands) according to the manufacturer’s protocol. For whole genome sequencing of single colonies, DNA from each colony was extracted using DNeasy Blood & Tissue Kit (Qiagen, Venlo, The Netherlands) according to the manufacturer’s protocol.

Libraries for metagenomic and whole genome sequencing were prepared using TruSeq Nano DNA (Illumina, San Diego, USA) with dual barcoded TruSeq DNA UD Indexes (Illumina, San Diego, USA). Sequencing was performed on an Illumina HiSeq X (Illumina, San Diego, USA) with v2.5 chemistry, producing 2 × 150 bp paired end reads. Metagenomic sequencing was performed in two separate runs.

### Recovery of Klebsiella pneumoniae

To recover *Klebsiella pneumoniae* from skim milk frozen stocks, 5 samples with *K. pneumoniae* MAGs were gradually defrosted on ice. 200 µL of the frozen stock were then added to 5 mL of BHI broth and incubated under shaking conditions (200 rpm) at 37°C overnight. The overnight culture was then serially diluted in PBS and 100 µL of the serially diluted culture was spread plated on Simmons citrate agar (Oxoid, Hampshire, UK) with inositol (Sigma-Aldrich, Missouri, USA) supplemented with 8 µg/mL of ampicillin (VWR, Pennsylvania, United States). After overnight incubation, 10 colonies were isolated from two samples (Sw1-Pier-TSA and Sw1-Pier-TSB), while 5 colonies were isolated from the other three samples (Sw1-Lag-BPW, Sw1-Lag-MacB, and Sw1-Lag-TSB). Each colony was then incubated overnight under shaking conditions, 200 rpm at 37°C in BHI broth and underwent rounds of purification. Prior to sequencing, colony PCR was performed using primers Pf (5′-ATTTGAAGAGGTTGCAAACGAT-3′) / Pr1 (5′-TTCACTCTGAAGTTTTCTTGTGTTC-3′) (Liu et al., 2008). The colony was isolated and suspended in nuclease-free water. The suspension was then boiled at 95°C for 10 min. 2 μL of supernatant was used as template DNA after centrifuge at 10,000 rpm for 5 min. For the PCR reaction, 2 μL template DNA was amplified in a 25 μL containing 1X Taq buffer with KCl, 50 mM KCl, 1.5 mM MgCl_2_, 0.1 mM each of the four dNTPs, 1-unit Taq DNA polymerase (Thermo Fisher Scientific, Massachusetts, USA), 1 μM of each primer and nuclease free water. The cycling conditions were 10 mins at 94°C followed by 35 cycles of 30 s at 94°C, 20 s at 57°C, and 20 s at 72°C then 10 mins hold at 72°C.

### Bioinformatics analysis

Kneaddata pipeline v0.7.7 (https://github.com/biobakery/kneaddata) was first used to remove low-quality reads and human contamination. Thereafter, draft taxonomy classification and relative abundance estimation were performed using MetaPhlAn4 with database: vJan21_CHOCOPhlAnSGB_202103 (Blanco-Míguez et al., 2023). Initial assembly of the contigs was generated using metaSPAdes v3.15.5 (Nurk et al., 2017). For binning, bin refinement and bin reassembly (threshold for bins was set as >50% completeness and <10% contamination rate), metaWRAP pipeline v1.3 was utilised (Uritskiy et al., 2018). For pure isolate genomes, Trimmomatic 0.39 and SKESA 2.5.1 within Shovill were utilised for quality control and removal of adapters as well as assembly, respectively (Seemann, 2016). For accurate taxonomic identification of the bins, the Genome Taxonomy Database (GTDB) and the associated taxonomic classification toolkit (GTDB-Tk) classify workflow was used (v2.1.1) using the Genome Taxonomy Database release 214 (Chaumeil et al., 2022). The bins and pure isolate genomes were annotated using Prokka v1.14.5 to predict protein-coding sequences (CDS) and RNA (Seemann, 2014). The detection of AMR genes in the MAGs was done using Staramr (Bharat et al., 2022). Core genome identification and alignment were performed using Roary v3.11.2 for the MAGs in this study and the reference genomes from the NCBI RefSeq database (Page et al., 2015). Maximum-likelihood trees were then built with RAxML v8.2.13 (Stamatakis, 2014) using the GTR□+□Gamma model with 1,000 times bootstrap based on the aligned core genes DNA sequences. For all software, default parameters were selected unless otherwise specified.

### Statistical analysis

Statistical analyses were performed in R version 4.3.3. Given that metagenomic relative abundance data are compositional and typically non-normally distributed, non-parametric tests were employed for all comparisons. A *p*-value of <0.05 was considered statistically significant. To evaluate the effect of different culture media (BPW, MacA, MacB, PseuA, TSA, and TSB) on the recovery of *Vibrio* spp. and *V. fluvialis* (with highest abundance), a Kruskal-Wallis rank-sum test was used. For significant results, post-hoc pairwise comparisons were conducted using Dunn’s test. To control the family-wise error rate associated with multiple comparisons, *p*-values were adjusted using the Benjamini-Hochberg (False Discovery Rate) method. For aggregate analysis, media were categorised into ‘Solid’ (PseuA, MacA, and TSA) and ‘Liquid’ (BPW, MacB, and TSB). The impact of the physical state of the media (Liquid vs. Solid) on microbial recovery was assessed using the Wilcoxon rank-sum test (Mann-Whitney U test). Data manipulation and visualization were conducted using the *tidyverse* suite (v2.0.0). Specific statistical tests were performed using the *stats* package and the *FSA* package (v0.9.5) for the Dunn’s test.

## Results and discussion

### Direct metagenomics show limited recovery of low-abundance pathogens in coastal waters

Across all sites, direct metagenomics of coastal water detected only abundant marine bacteria, with only 0.03% of sequencing reads from human pathogens based on a total sequencing output of 11.50 Gbp (**Figure *3***). In contrast, culture-enrichment using various media markedly increased the proportion (21.2%) of sequencing reads corresponding to low-abundance ESKAPE pathogens and pathogenic *Vibrio* spp., with a total sequencing output of 275.2 Gbp for 33 enriched coastal water samples. It represents a 700-fold increase in relative abundance compared to direct metagenomic samples, despite the higher sequencing depth employed for the enriched library. The result is consistent with a previous study where Enterobacterales represented less than 1% of direct metagenomics sequencing reads but comprised between 15 – 20% of enriched metagenomics of freshwater samples (Ottesen et al., 2022). The results also highlight that culture-enrichment effectively enriches clinically relevant taxa, although recovery is biased toward organisms suited to each medium, direct metagenomics entirely misses these low-abundance but important pathogens. Pathogens were defined as bacteria capable of causing human infection, based on the list compiled by Bartlett et al. (Bartlett et al., 2022). The medium used in culture-enrichment were Buffered Peptone Water (BPW), MacConkey broth (MacB), tryptic soy broth (TSB), Pseudomonas agar (PseuA), MacConkey agar (MacA), and tryptic soy agar (TSA).

**Figure 3.**
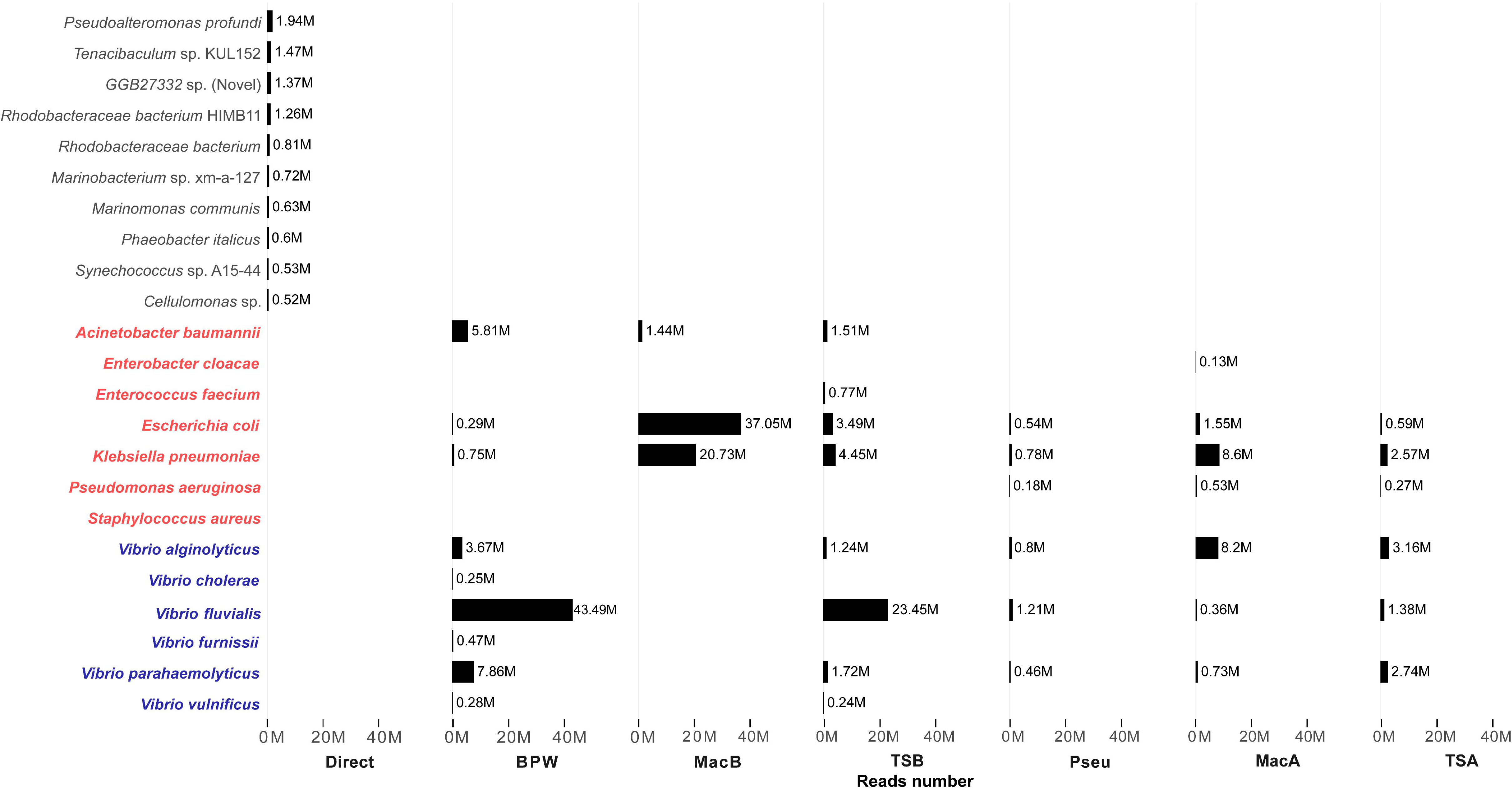
Number of sequencing reads obtained from direct (n=6) and culture-enriched metagenomics of all coastal water samples (n=6 for BPW, MacB, TSB, MacA, n=5 for TSA and n=4 for PseuA). The top 10 species detected by direct metagenomics, ESKAPE pathogens (*Acinetobacter baumannii*, *Enterobacter cloacae*, *Enterococcus faecium*, *Escherichia coli*, *Klebsiella pneumoniae*, *Pseudomonas aeruginosa*, and *Staphylococcus aureus*), and pathogenic *Vibrio* spp. are shown. Only read counts greater than 0.1 million (0.1M) are displayed. ESKAPE pathogens are highlighted in red, and pathogenic *Vibrio* spp. are highlighted in blue. Pathogens were classified based on the human pathogen list reported by Bartlett et al. (Bartlett et al., 2022). Abbreviations: BPW, Buffered peptone water; MacB, MacConkey broth; TSB, Tryptic soy broth; PseuA, Pseudomonas agar; MacA, MacConkey agar; TSA, Tryptic soy agar.

To allow direct comparison between libraries of different sequencing output, the abundance of pathogens was normalised to reads per million (RPM), calculated as number of reads assigned to a species divide by total library reads for respective enrichment medium and multiply by 10^6^. While direct metagenomics captured only a negligible pathogen signal (combined 300 RPM for all target species), culture-enriched metagenomics markedly concentrated ESKAPE and *Vibrio* spp. across all water samples. Among the media tested, PseuA had the lowest combined pathogen density (42,275 RPM), while MacB achieved the highest enrichment (334,028 RPM) for target species. This represents at least 140-fold increase in detection frequency, independent of sequencing depth (**Supplementary table S1**). Notably, several ESKAPE species were only detectable (RPM > 0) following culture-enrichment.

Culture-enriched metagenomics of the six coastal water samples showed that for ESKAPE pathogens, reads corresponding to *Acinetobacter baumannii* were detected only in the liquid media BPW (2.9%), MacB (0.81%), and TSB (0.91%) (the value in parentheses represents the percentage of the reads relative to the total number of reads generated from all samples combined within each respective enrichment medium). Conversely, *Pseudomonas aeruginosa* reads were recovered only from solid media, namely PseuA (0.19%), MacA (0.36%), and TSA (0.21%). For *Enterobacter cloacae* and *Enterococcus faecium*, recovery was observed only in MacA (0.1%) and TSB (0.46%), respectively. No reads corresponding to *Staphylococcus aureus* were detected in any of the culture enrichment conditions for water samples. All culture media were able to recover *Klebsiella pneumoniae* and *Escherichia coli*, albeit to different extents. As expected, MacConkey broth (MacB), a selective media for Enterobacterales, showed superior recovery of *K. pneumoniae* and *E. coli* (20.84% for *E. coli* and 11.65% for *K. pneumoniae*). *K. pneumoniae* and *E. coli* were also able to grow in generic media such as TSB (2.1% for *E. coli* and 2.67% for *K. pneumoniae*).

Among the media tested, BPW in particular recovered reads of the six main pathogenic *Vibrio* spp.: *V. alginolyticus*, *V. cholerae*, *V. fluvialis*, *V. furnissii*, *V. parahaemolyticus* and *V. vulnificus*. *V. fluvialis* had the highest read counts across the culture enrichment conditions, contributed mostly by BPW (21.67%) and TSB (14.08%). *V. parahaemolyticus* were also recovered at high abundances from the water samples by BPW (3.92%) and TSA (2.14%). Coincidentally, *V. parahaemolyticus* and *V. fluvialis* have the highest incidence rates among all *Vibrio* spp. infections in Singapore (Neoh et al., 2026). No reads corresponding to pathogenic *Vibrio* spp. was detected in MacB, indicating MacConkey media specificity towards gram-negative enteric bacteria (Allen, 2005). MacA recovered some *Vibrio* spp., specifically *V. alginolyticus* (5.63%), *V. parahaemolyticus* (0.5%) and *V. fluvialis* (0.25%). Although APW is usually preferred for growing *Vibrio* spp., our results show that BPW is also effective in enriching pathogenic members of this genus. (Centers for Disease Control and Prevention (CDC), 2024)

In terms of alpha diversity (**Supplementary figure S1)**, direct metagenomics of coastal water unsurprisingly exhibited the highest values for Observed Species, Shannon, and Simpson indices, capturing the most complete and even representation of the seawater microbiome. Non-selective TSB and BPW retained relatively high richness and evenness, particularly in water samples, suggesting these media capture a significant part of the community diversity. In contrast, selective MacA and MacB enrichments displayed lowest alpha diversity, especially within sediment and seaweed samples, where richness was markedly reduced. This indicated a strong selective bottleneck in these media that favours specific taxa over the broader community. TSA and PseuA recovered communities with moderate richness and evenness that were not dominated by a single taxon.

The rank-abundance comparison between direct and culture-enriched metagenomics revealed that culture-enriched methods strongly enhanced the detection of ESKAPE pathogens, especially when these are present at very low abundance (**Supplementary Figure S2**), highlighting the utility of culture-enriched metagenomics in effective pathogen surveillance. Among sample types, water samples exhibited the highest ESKAPE diversity, whereas seaweed showed the lowest, suggesting host selection. MacB outperformed other media in terms of efficiency and selectivity, yielding ESKAPE species as the dominant taxa with high relative abundance.

### Culture-enriched metagenomics clearly identifies subtle bacterial signatures of niche selection and human/animal impact

Bray-Curtis distance (**Figure *4***) is a useful metric to quantify the dissimilarity between microbial communities across samples. While each culture-enrichment medium recovered a distinct environmental microbiome profile, all media consistently yielded higher abundances of pathogenic bacteria for the different environmental matrices. Notably, *Staphylococcus* spp., specifically *S. warneri* and *S. epidermidis*, were detected only in solid media (TSA and MacA) (**Figure *4*A**). Gram positive *Bacilli* as well as most of the *Vibrio spp.* also favoured solid media. Unsurprisingly, enteric bacteria such as *Klebsiella pneumoniae*, *Escherichia coli*, *Enterobacter hormachei*, *Klebsiella aerogenes* and *Aeromonas hydrophila* preferentially grew on MacConkey media. Aquatic bacterial pathogens such as *Vibrio parahaemolyticus*, *V. fluvialis*, and *Shewanella algae* were recovered at higher abundance in BPW, while *Acinetobacter baumannii* and *Enterococcus faecalis* were more frequently enriched in TSB. PseuA did not recover *Pseudomonas* spp. optimally as expected, while it recovered *Pantoea anthophila* and *Citrobacter amalonaticus* at high abundances. In contrast with enrichments being dominated by human pathogens, direct metagenomics was dominated by marine Alphaproteobacteria.

**Figure 4.**
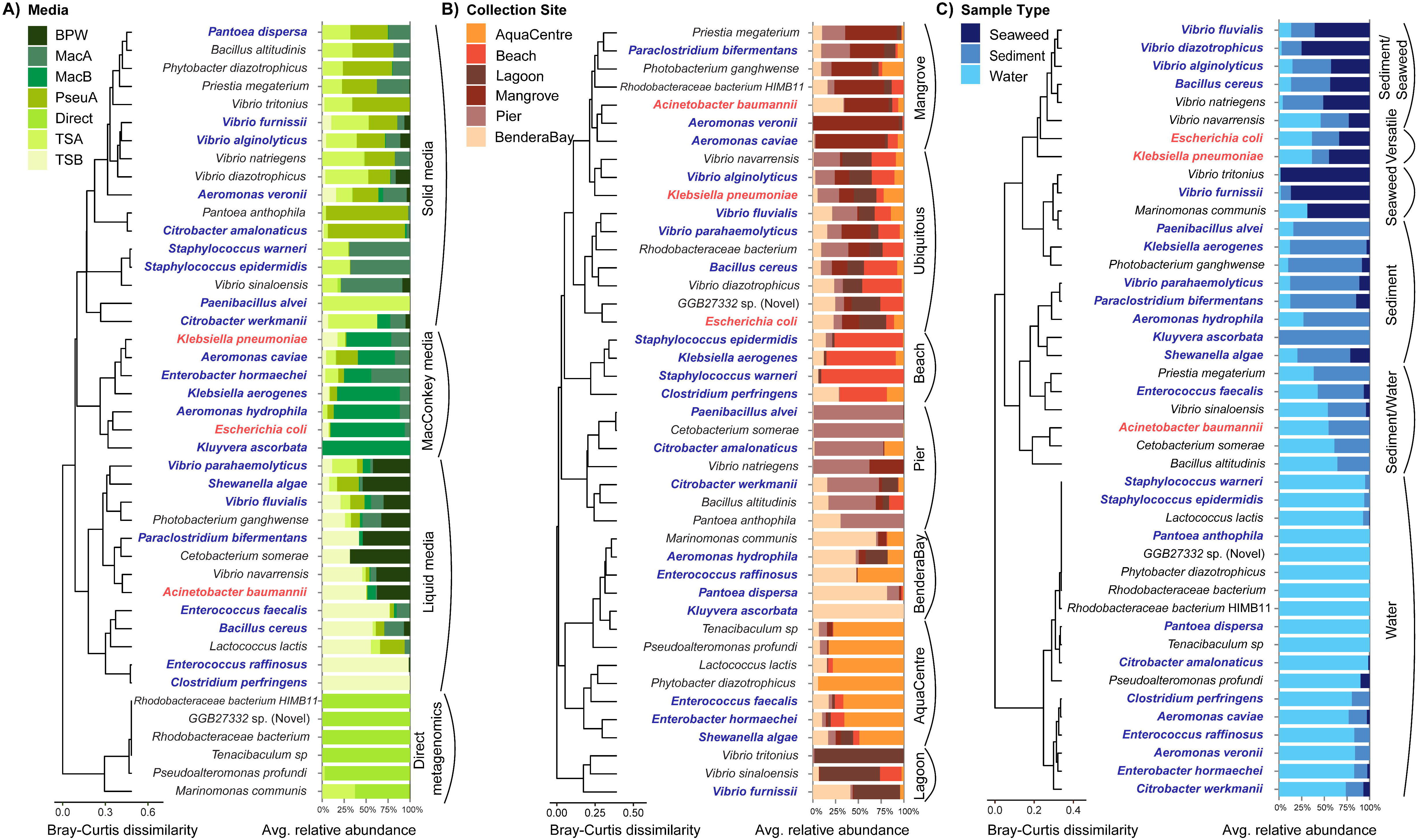
Bray-Curtis clustering based on species-level relative abundance profiles of direct metagenomics and culture-enriched metagenomics of all three sample matrices. Profiles are grouped by (A) culture media, (B) collection site, and (C) sample type. Bar plots show the average relative abundance of each species within each category. ESKAPE pathogens are highlighted in red, and other pathogens are highlighted in blue. Pathogens were classified based on the human pathogen list reported by Bartlett et al. (Bartlett et al., 2022). Abbreviations: BPW, Buffered peptone water; MacB, MacConkey broth; TSB, Tryptic soy broth; PseuA, Pseudomonas agar; MacA, MacConkey agar; TSA, Tryptic soy agar, AquaCentre: Aquaculture centre.

By employing culture-enriched metagenomics, site-specific microbial signatures were observed across the sampling locations (**Figure *4*B**). At the sampling site near the outlet of the aquaculture centre (AquaCentre), bacteria such as *E. faecalis* and *S. algae* were present at higher abundance, as well as the fish-associated pathogen, *Tenacibaculum* spp. The latter causes high mortalities for marine fishes through an ulcerative disease, tenacibaculosis (Mabrok et al., 2022, Miyake et al., 2020). Bendera Bay, which is a restricted area with few visitors, exhibited a distinct microbiome profile compared with the other sampling sites with a higher abundance of *Aeromonas hydrophila*, *Enterococcus raffinosus*, *Pantoea dispersa* and *Kluyvera ascorbata*. These species represent a group of emerging opportunistic pathogens with a high degree of environmental plasticity. For instance, *A. hydrophila* is a known causative agent of diarrheal disease while *E. raffinosus* and *P. dispersa* are increasingly associated and notorious for causing systemic and nosocomial infections, such as bacteremia (Pessoa et al., 2022, Panditrao and Panditrao, 2018, Lee et al., 2022). Notably, both *E. raffinosus* and *K. ascorbata* serve as significant reservoirs for AMR, facilitating the dissemination of resistance genes across diverse ecological niches (Wilke et al., 1997, Rautela et al., 2024, Artunduaga-Cañas et al., 2025). Presence of these bacteria further highlights the site’s potential as a hotspot for horizontal gene transfer of AMR. Human-associated bacteria, including *S. epidermidis*, *K. aerogenes*, *S. warneri*, and *Clostridium perfringens*, were detected at higher abundance in area where human traffic is intense, such as the beach. In addition, pathogenic bacteria such as *E. coli*, *K. pneumoniae*, *V. parahaemolyticus*, *V. fluvialis*, and *V. alginolyticus* were ubiquitously detected across all collection sites, suggesting environmental resilience and potential for dissemination (Brumfield et al., 2023). Their presence is increasingly linked to climate change, with warmer water temperatures facilitating their proliferation (Lee et al., 2019). In contrast, samples from the mangrove patch showed a microbiome profile enriched in *A. baumannii*, *Aeromonas veronii*, and *A. caviae*. At the pier, several carbon-degrading species, including *Paenibacillus alvei*, *C. amalonaticus*, *C. werkmanii*, *Bacillus altitudinis*, *P. anthopila* were present alongside *Cetobacterium somerae*. *C. somerae* is a predominant bacterium within the intestinal microbiota of fish, correlating with the schools of fish observed near the ferry berthing area (Maervoet et al., 2012, Ogunbayo et al., 2018, Grady et al., 2016, Tsuchiya et al., 2008). *Cetobacterium* spp. was also shown to be part of gut microbiota among farmed Asian seabass in Singapore (Soh et al., 2025). None of these signatures could be observed by direct metagenomics.

Culture-enriched metagenomics enabled the detection of pathogen-specific distributions, with certain lineages occurring more frequently in seaweed, sediment, or water samples (**Figure *4*C**). *E. coli*, *K. pneumoniae* and *Vibrio navarrensis* were detected across all sample types and geographically widespread on the island. *A. baumannii* and *E. faecalis* were more abundant in water and sediment samples, suggesting exclusion by macroalgal hosts. Pathogenic *Vibrio* spp. were more commonly detected on seaweed, except for *V. parahaemolyticus*, which was mostly found in sediment. This observation aligns with the known ecology of *V. parahaemolyticus*, which resides in the benthos before being resuspended into the water column in correlation with rising water temperatures (Johnson et al., 2010, Su and Liu, 2007). Some species were almost exclusively found in the water, including *S. warneri*, *S. epidermidis*, *Lactococcus* spp. and Alphaproteobacteria. *S. algae* were more abundant in sediment but were also detected in water and seaweed samples. Previous studies have identified *S. algae* in coastal water and oysters in Taiwan and the US (Tseng et al., 2018, Johnson et al., 2025), but this facultative anaerobe can metabolise various inorganic and organic compounds and can exist at high abundance in coastal sediments (Venkateswaran et al., 1999, Inohana et al., 2020, Shen et al., 2025). It also tolerates a wide range of temperature and salinities (Tseng et al., 2018). These characteristics highlight the need for environmental surveillance of *S. algae*; consequently, we characterised *S. algae* MAGs from this study and their associated AMR profiles in greater detail elsewhere (Ho et al., 2025a).

### Culture-enriched metagenomics can rapidly identify biases of growth media

The ESKAPE genera were all present in the top 20 organisms recovered by culture-enriched metagenomics (**Figure *5*A)**, showing that human pathogens are preferentially amplified over non-pathogenic species through enrichment. The *Vibrio* genus had the highest mean relative abundance recovered by all the media used, with solid (38.9%) and liquid (37.3%) media having similar recovery rates, and there is no significant difference observed between liquid and solid media overall (Wilcoxon rank-sum, W=931, p = 0.80). The percentages represent the mean relative abundance of each genus, calculated as the average relative abundance across all samples within each media group. There was a significant difference in the relative abundance of *Vibrio* spp. across the six media tested (Kruskal-Wallis, X^2^ = 19.44, df = 5, p = 0.002). Post-hoc pairwise comparisons using Dunn’s test (with Benjamini-Hochberg adjustment) specifically identified that BPW achieved significantly higher relative abundance of *Vibrio* spp. (59.11%) compared to MacB (p < 0.001), of which *V. fluvialis* accounted for a 37.63% share. While MacB consistently showed lower mean recovery (15.48%) than all other media tested for *Vibrio* spp., these differences were only statistically significant when compared to BPW. **Figure *5*B** shows the mean relative abundance of the five main *Vibrio* pathogenic species in enrichments from all sample matrices: *V. cholera*, *V. fluvialis*, *V. furnissii*, *V. parahaemolyticus* and *V. vulnificus,* as well as other minor pathogenic species. *V. fluvialis* was efficiently recovered with most media (19.26% - 37.63%), except MacB (11.86%). The recovery of *V. fluvialis* was significantly affected by the choice of culture media (Kruskal-Wallis, p = 0.0075). Post-hoc analysis confirmed that BPW yielded a significantly higher relative abundance of *V. fluvialis* compared to MacB (Dunn’s test, p = 0.004). *V. parahaemolyticus* had the second highest mean relative abundance among them, which is also reflected in the high number of MAGs recovered (**Table *1***). *V. parahaemolyticus* had been detected carrying OXA-48-like ARGs in the region, hence requiring surveillance due to remarkable genetic plasticity in *Vibrio* spp. (Goh et al., 2022, Yuan et al., 2025). All the media used had limited recovery for other pathogenic *Vibrio* spp., *V. cholerae* recording relative abundance of <0.2%, while *V. mimicus* and *V. metoecus* were <0.1%. Although abundance of both *V. cholerae* and *V. metoecus* is known to be lower in marine environments as opposed to brackish or freshwater (Kirchberger et al., 2014), the presence of *V. cholerae* in Singapore coastal water has been shown by other study (Goh et al., 2017). Our results demonstrated that specific nutrient composition and selective agents, rather than the physical state of the media (liquid or solid), dictate recovery success for coastal *Vibrio* species. This further reinforces that bacteria such as *V. cholerae* require specific enrichment media, such as thiosulfate citrate bile-salts sucrose (TCBS) agar, tellurite taurocholate gelatin agar (TTGA), and CHROMagar™ Vibrio (Huq et al., 2012), while this is not the case for *V. fluvialis* or *V. parahaemolyticus*. *V. cholerae* could also enter into a viable but nonculturable (VBNC) state in which they may not form colonies on traditional bacteriological culture plates (Roszak and Colwell, 1987).

**Figure 5.**
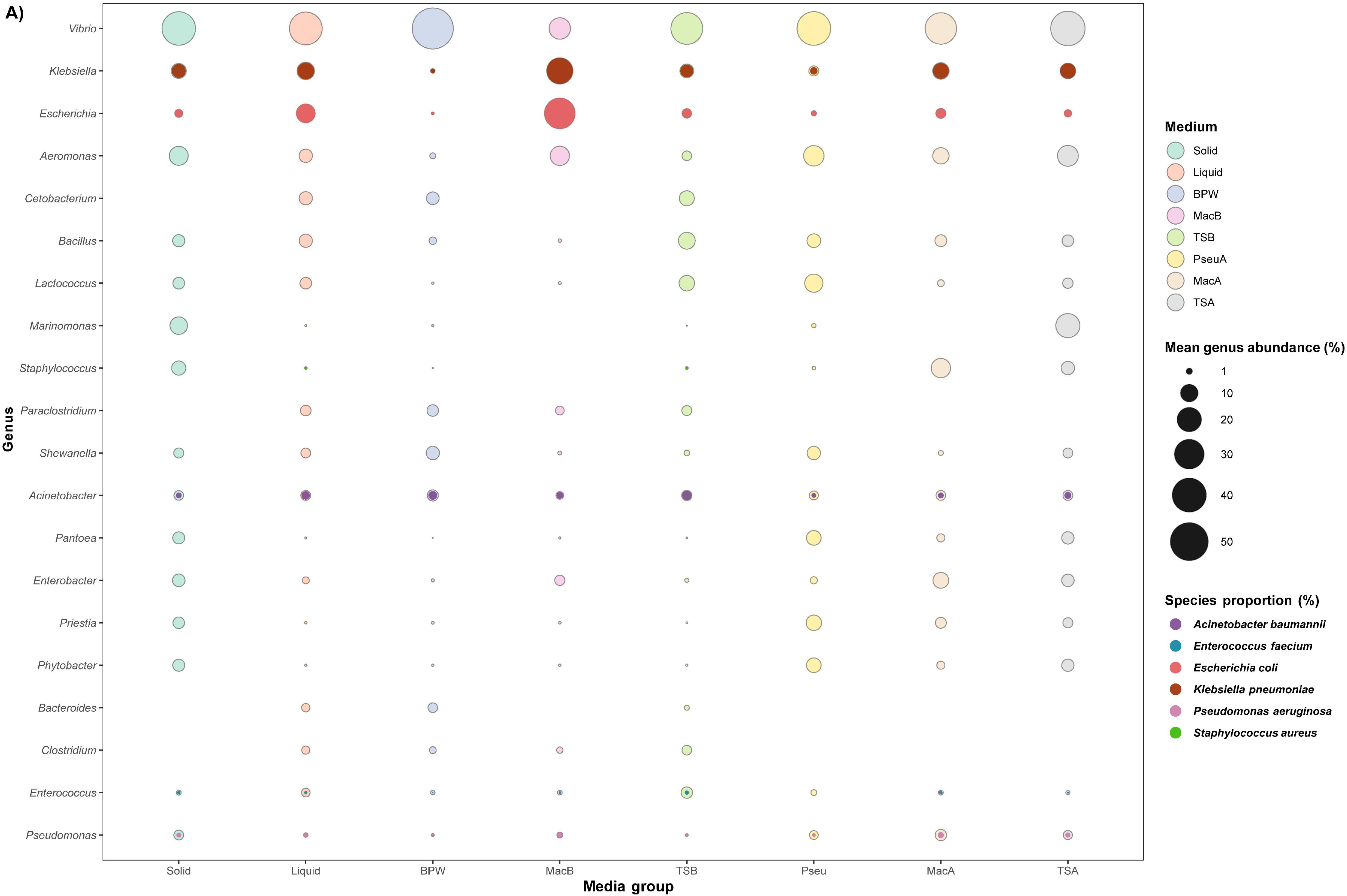
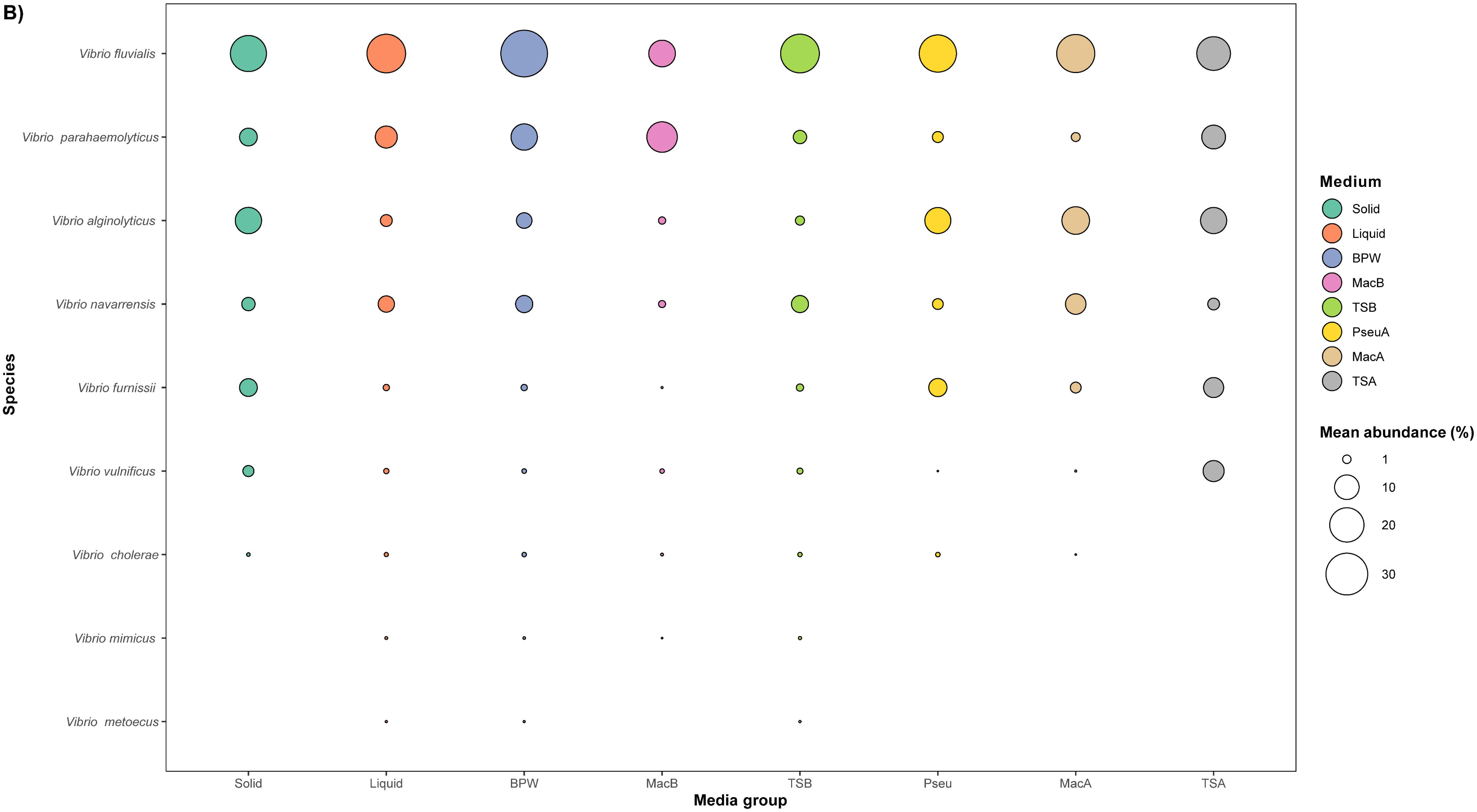
Overall distribution of dominant taxa across culture-enrichment conditions. Colours represent different media; solid media include PseuA, MacA, and TSA, while liquid media include BPW, MacB, and TSB. (A) Top 20 genera with the highest relative abundance across all culture-enrichment conditions (in descending order). Outer circles with lower opacity represent the mean genus-level relative abundance. Inner circles indicate the proportional contribution of target ESKAPE species within each genus, scaled to the corresponding genus abundance. (B) Relative abundance and distribution of *Vibrio* spp. (*V. fluvialis*, *V. parahaemolyticus*, *V. alginolyticus*, *V. navarrensis*, *V. furnissii*, *V. vulnificus*, *V. cholerae*, *V. mimicus*, and *V. metoecus*) across culture-enrichment conditions (in descending order). Circle size represents the mean relative abundance of each species. Abbreviations: BPW, Buffered peptone water; MacB, MacConkey broth; TSB, Tryptic soy broth; PseuA, Pseudomonas agar; MacA, MacConkey agar; TSA, Tryptic soy agar.

**Table 1.**
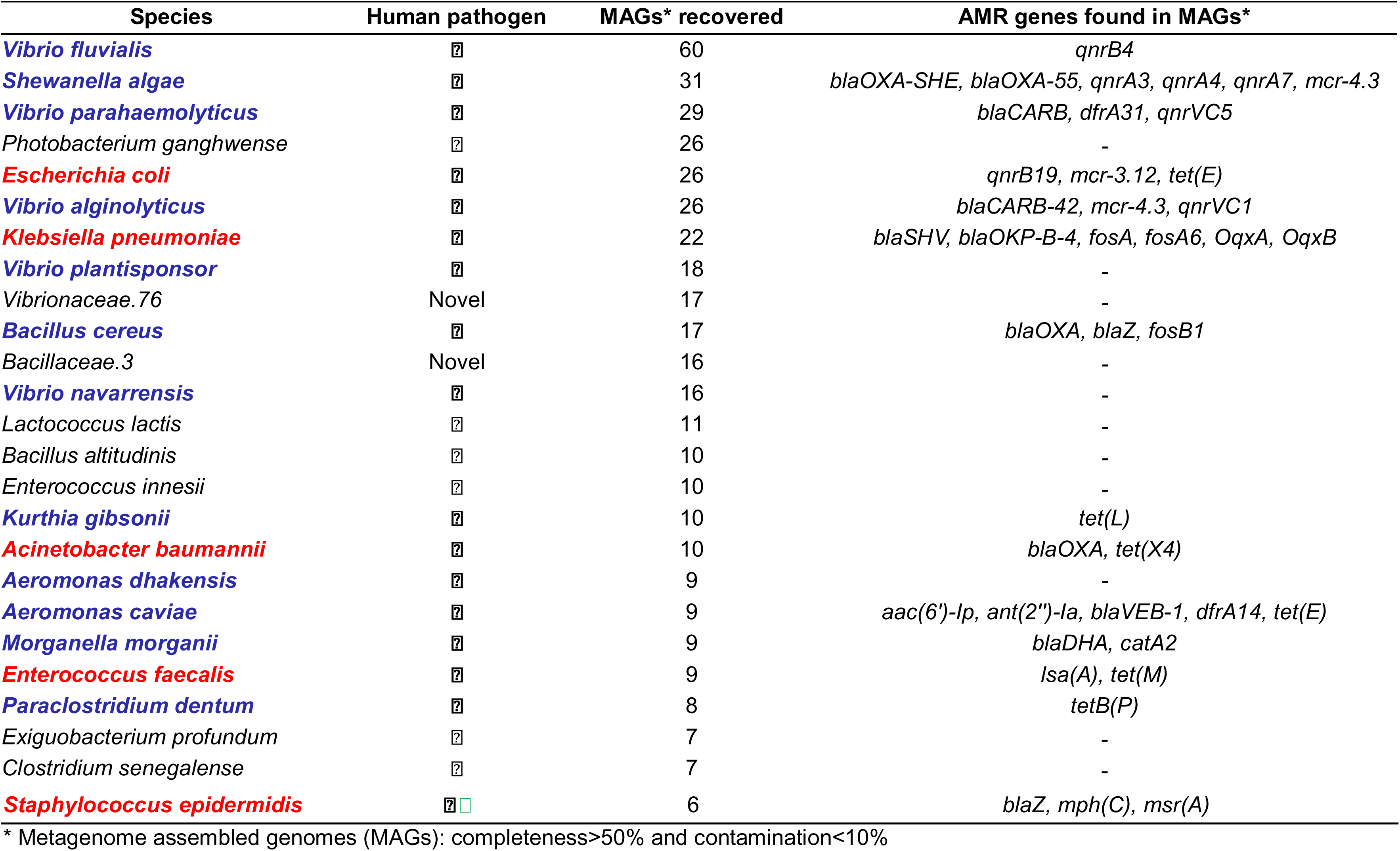
Top 25 species for which genomes were recovered by culture-enriched metagenomics and its associated AMR genes. ESKAPE pathogens are highlighted in red, and other pathogens are highlighted in blue. Pathogens were classified based on the human pathogen list reported by Bartlett et al. (Bartlett et al., 2022). The detection of AMR genes in the MAGs was done using Staramr (Bharat et al., 2022).

*Klebsiella* and *Escherichia* were among the top three most abundant genera recovered from culture-enriched metagenomics, and most of the species recovered were the top priority pathogens, *K. pneumoniae* and *E. coli* (**Figure *5*A**). For the focus species, (*E. coli*, *S. aureus, K. pneumoniae*, *A. baumannii*, *P. aeruginosa,* and *E. faecium*), proportions were calculated by dividing the relative abundance of the species by the total relative abundance of its parent genus. These values were then averaged across all samples for each media group and represented as percentages. Liquid media, specifically MacB, outperformed the solid media in recovering both *Klebsiella* and *Escherichia* spp., including *K. pneumoniae* and *E. coli*. Almost all of the *Escherichia* spp. recovered using culture-enrichment (all >99.5%) were *E. coli*. For *Staphylococcus* spp., there was a significant bias towards solid media, such as MacA and TSA (Wilcoxon rank-sum, W = 31, p = 0.005). However, only TSB liquid media could recover *S. aureus*. This indicates that *S. aureus* in particular preferred liquid media, but that this is not the case for all *Staphylococcus* spp. in general (Missiakas and Schneewind, 2013). At the genus level, *Acinetobacter* spp. had similar average relative abundances for both solid and liquid enrichment (Solid: 2.8%; Liquid: 2.95%). In contrast, the recovery proportion of *Acinetobacter* spp. belonging to the *A. baumannii* species was higher in liquid media (68.27%) than in solid media (29.43%). For *Enterococcus* spp., the average relative abundance was low in both type of enrichment (Solid: 0.6%, Liquid: 2.17%). At the genus level, *Pseudomonas* spp. represented a mean relative abundance of 2.9% in solid media compared to 0.46% in liquid media, and the difference is statistically significant (Wilcoxon rank sum, W = 78, p < 0.001). However, within this genus, *P. aeruginosa* accounted for a higher intra-genus proportion in liquid media (58.09%) largely driven by its recovery in MacB, where it comprised 82.35% of all *Pseudomonas* spp. Some of the genera were biased towards liquid enrichment for recovery, including *Cetobacterium*, *Paraclostridium*, *Bacteroides* and *Clostridium*. Growth for these taxa was exclusively observed in liquid enrichment, with no detectable relative abundance in solid media across all sample matrices.

### Culture-enriched metagenomics enables streamlined and cost-effective recovery of pathogen genomes

While total microbial diversity decreased in enrichment cultures due to selective pressure exerted by the media, the recovery of medium to high-quality MAGs (≥50% completeness, <10% contamination) increased substantially. The pipeline yielded 1,124 MAGs in total. Enrichment samples (n = 92; 800.49 Gbp) contributed 773 MAGs, whereas direct coastal water sequencing (n = 6; 11.50 Gbp) yielded 351 MAGs. Among these, 427 MAGs recovered from culture-enriched metagenomics and 117 MAGs from direct metagenomics were high quality MAGs (>90% completeness and <5% contamination).

However, at any taxonomic level, direct metagenomics yielded a higher proportion (307 MAGs) of unidentified MAGs, lacking taxonomic classification, whereas only 163 unidentified MAGs were recovered across all enrichment samples from different matrices. In direct metagenomics, only 44 MAGs were identified at the species level, and 16 were identified only to the genus level. Among the 44 species-level MAGs, 42 were from non-pathogenic species, with only two exceptions: *Kytococcus sedentarius* and *Dietza maris.* Both species are opportunistic pathogens of environmental origin, although human infection by these two organisms is relatively uncommon (Ioannou et al., 2025, Sims et al., 2009, Tamarit et al., 2007, Perkin et al., 2012). In contrast, enrichment samples produced 113 MAGs that were unidentified at the species level and 610 with a species identification. Among the latter, 383 MAGs (62.8%) belonged to known pathogenic species and 227 to non-pathogens. The top 25 most frequently recovered MAGs were from known pathogenic species, including *V. fluvialis, S. algae, V. parahaemolyticus, E. coli, V. alginolyticus, K. pneumoniae, A. baumannii, E. faecalis and S. epidermidis* (Table 1). All of these MAGs corresponding from pathogenic species were recovered exclusively from culture-enriched metagenomics, with none obtained from direct metagenomics. This shows that while culture-enrichment involved additional media costs, it significantly reduces the total sequencing depth required to recover low-abundance pathogen genomes that was entirely unable to be recovered by direct metagenomics. Specifically, given that the target pathogen density in MacB enrichment of water samples (334,028 RPM) was 1,100 times higher than in direct metagenomics (300 RPM) (**Figure *3*** and **Supplementary table S1**), achieving equivalent genomic resolution through direct metagenomics would require a cost increase of approximately three orders of magnitude. Thus, culture-enrichment offers a financially viable pathway for routine environmental AMR surveillance, particularly in resource-limited settings where ultra-deep sequencing is not feasible. The recovered MAGs were also of sufficiently high quality to allow the detection of antimicrobial resistance (AMR) genes associated with them (**Supplementary table S2**). Notably, one of our *V. parahaemolyticus* MAGs is genetically highly similar to a local hospital isolate (**Supplementary figure S3**), with multilocus sequence typing (MLST) analysis indicating that it is a single locus variant belonging to the same clonal complex (**Supplementary table S3, S4**). Haemolytic AMR *V. parahaemolyticus* had previously been detected in farmed green mussels and farm water in Singapore (Ong et al., 2023). This raises concern of food safety, as human diarrhoeal diseases linked to seafood consumption are mostly caused by *V. parahaemolyticus* an *V. vulnificus* (Muzembo et al., 2024). In Southeast Asia, despite being underreported, *V. parahaemolyticus* has been identified as the primary cause of non-cholera vibriosis in diarrheal patients (Muzembo et al., 2024, Neoh et al., 2026).

To validate the MAGs recovered from culture-enriched metagenomics, we isolated *K. pneumoniae* colonies from preserved enrichment stocks and performed whole-genome sequencing. The assembled genomes from our culture-enriched metagenomics showed high degree of genotypic congruence to the genomes from pure isolates (**Figure *6***), confirming their suitability for epidemiological studies. In the phylogenetic tree constructed, MAGs highlighted the presence of multidrug-resistant lineages, many of which matching globally prevalent clones. For example, MAGs SJIM33.5 and SJIM 11.8 were closely related to three isolates SJI33, 35 and 37, with the same ST716 identified. Recently, the ST716 lineage was identified as the cause of a carbapenem-resistant strain outbreak within a neurorehabilitation unit (Silvotti et al., 2025). Genome annotations revealed the co-occurrence of multiple resistance and virulence determinants, raising concerns about environmental reservoirs of clinically important strains. While WGS of pure isolates remains the gold standard for genomic accuracy, our culture-enriched *K. pneumoniae* MAGs achieved comparable quality and allowed high strain-level resolution, to the clonal complex level as defined in MLST (**Supplementary table S5)**. Of the 22 *K. pneumoniae* MAGs, 12 (54.5%) were of high quality (>90% completeness, <5% contamination) and the rest were of medium quality (≥50% completeness, <10% contamination). Qualitatively, the MAGs captured a broader spectrum of diversity that might be overlooked during manual colony picking (**Figure *6***). While the WGS isolates provided a more definitive genomic scaffold required to validate these MAGs, our MAGs recovered from culture-enriched metagenomics also captured a comparable breadth of genomic features. The MAGs maintained the integrity of key epidemiological markers, such as MLST Sequence Type (ST), O- and K-antigen profiles, matching the resolution of pure isolates. **Supplementary table S5** highlights the close genotypic alignment between environmental MAGs, pure isolates and clinical reference strains for *K. pneumoniae*. For instance, we identified identical STs and shared ARG profiles across different sources, such as the ST716 and the *bla*_SHV-27_ gene found in our MAGs, isolates, and international reference strains. This indicates the accuracy of MAG recovery from culture-enriched metagenomics in reflecting clinically relevant resistance patterns. It also demonstrates that culture-enriched methods support the effectiveness of targeted culturing for pathogen detection in environmental samples, a critical consideration given that pandemic *E. coli* strains in Singapore show strong genotypic similarity between environmental and hospital-derived isolates (Yuan et al., 2024).

**Figure 6.**
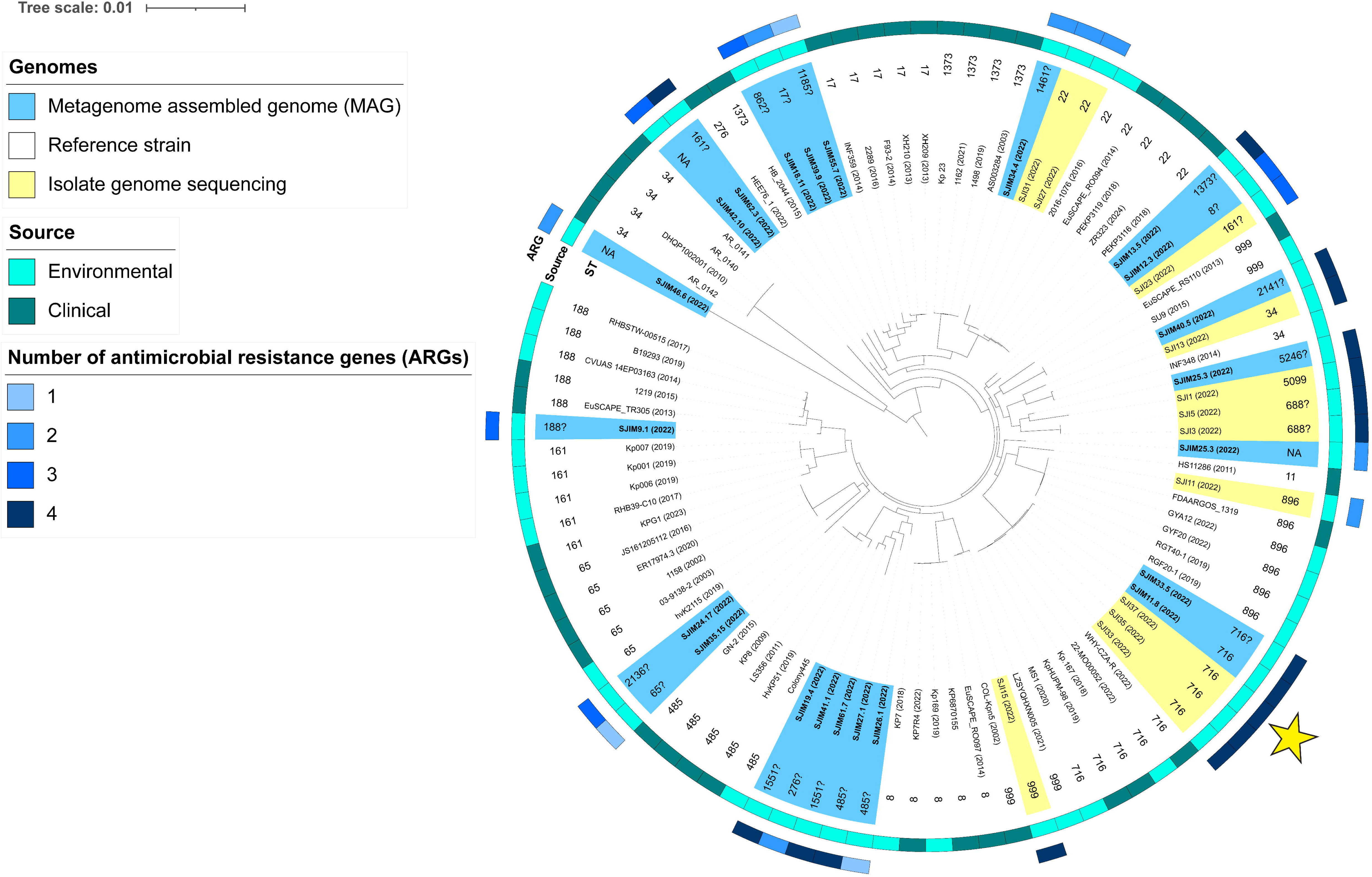
Circular core-genome maximum-likelihood trees of *Klebsiella pneumoniae*. (From innermost to outermost) The colour at the first ring denotes the genome types, including MAGs, reference strains and isolate genome sequencing. The second ring indicates the Sequence Types (ST) types of the genome, while the third ring indicates the sources of the genomes whether from the environment or clinical. The outermost ring indicates number of antimicrobial resistance genes (ARGs) detected in the MAGs and isolate genomes from this study in different colours. The core genome tree is rooted to *K. quasipneumoniae* with 478 core genes aligned. The scale bar indicates 0.01 nucleotide per site. Yellow star indicates where MAGs SJIM33.5 and SJIM 11.8 were closely related to three isolates SJI33, 35 and 37, with the same ST716 identified. ST followed by a question mark (?) represent presumptive assignments based on the highest probability match. These occur where MAG incompleteness or minor sequence variations prevent a 100% identity match to the MLST schema, though the partial allelic profile remains most consistent with the labelled ST.

## 4 Conclusions

Our study demonstrates that culture-enriched metagenomics can recover high-quality genomes of low-abundance, clinically important bacteria from complex environmental samples and uncover the hidden diversity of environmental pathogens. This method holds promise for environmental AMR surveillance, particularly in high-risk zones such as tropical coasts with high anthropogenic pressure, including Singapore (Sin et al., 2016). In coastal habitats such as those around St. John’s Island, this method reveals the widespread presence of ESKAPE and *Vibrio* species, many with resistance traits, and provides a high-resolution framework for AMR risk assessment and microbial ecology. The presence of multidrug-resistant *K. pneumoniae* and other pathogens in sediments and seaweed-associated microbiomes points to their potential persistence and spread in marine ecosystems. Our results established culture-enriched metagenomics as a robust tool to uncover specific microbial signatures that serve as indicators of niche-specific selection and anthropogenic pressures. Future work should explore temporal trends, mobile genetic elements, and environmental drivers of AMR dissemination.

## Supplementary figure

**Supplementary figure S1 Alpha diversity across sample types (seaweed, sediment and water) for direct and culture-enriched metagenomics. The boxplot shows the distribution of alpha diversity value across the samples for each sample type (seaweed, n=4, sediment, n=6; water, n=6).**

Abbreviations: BPW, Buffered peptone water; MacB, MacConkey broth; TSB, Tryptic soy broth; PseuA, Pseudomonas agar; MacA, MacConkey agar; TSA, Tryptic soy agar.

**Supplementary figure S2 Rank abundance curves comparing microbial community structure between direct metagenomics (raw) and culture-enriched methods (BPW, MacA, MacB, PseuA, TSA and TSB) across seaweed (n=4), sediment (n=6) and water (n=6) samples. Taxa were rank ordered from highest to lowest abundance (left to right).**

**Supplementary figure S3 Maximum-likelihood tree of all *Vibrio parahaemolyticus* MAGs (except two with completeness of less than 80%) including four addition reference genomes in blue text and a strain from the hospital (TTSH S145) in red text. The core genome tree is mid-point rooted and is based on 1538 core genes. The scale bar indicates 0.001 nucleotide substitution per site.**

## Author contribution

Jia Yee Ho: Data curation; Formal analysis; Investigation; Methodology; Project administration; Software; Validation; Visualization; Writing - original draft; Writing - review & editing. Dalong Hu: Data curation; Formal analysis; Investigation; Methodology; Software; Validation; Visualization; Writing - review & editing. Deborah Yebon Kang: Investigation, Writing - review & editing. Clarence Bo Wen Sim: Investigation, Writing - review & editing. Winona Wijaya: Software; Validation; Visualization; Writing - review & editing. Yann Felix Boucher: Conceptualization; Funding acquisition; Methodology; Project administration; Resources; Supervision; Writing - review & editing.

## Acknowledgements

The Singapore Centre for Environmental Life Sciences Engineering (SCELSE) is funded by the Ministry of Education, Singapore, the National Research Foundation of Singapore, Nanyang Technological University Singapore (NTU) and National University of Singapore (NUS). We would like to thank the National University of Singapore Information Technology for their support in management of the high-performance computing, the St. John’s Island National Marine Laboratory for providing the facility necessary for conducting the research; the Laboratory is a National Research Infrastructure under the National Research Foundation Singapore. We would also like to acknowledge the National Parks Board (NParks) Singapore for granting the research permit at St. John’s Island and extend our gratitude to Mr. Sebastian Jie Hong Yeo for his immense help in the fieldwork. We would like to thank Dr. Yihui Chen from Tan Tock Seng Hospital for sharing *Vibrio parahaemolyticus* genomic data for comparative analysis.

## Data availability statement

This metagenomics sequencing project has been deposited in the DDBJ Sequence Read Archive (DRA) under BioProject PRJDB17097. The raw reads used for assembling the MAGs and MAGs are associated with BioSample numbers <u>SAMD00659505</u>–<u>SAMD00659561</u>, <u>SAMD00731966</u>–<u>SAMD00732000</u>, <u>SAMD00732048</u>–<u>SAMD00732053</u>, <u>SAMD01828761</u>–<u>SAMD01828811</u>, and <u>SAMD01838622</u>–<u>SAMD01838633</u>.

