## Supplementary figures and images for "Culture-enriched metagenomics enables genome-resolved detection of low abundance ESKAPE and *Vibrio* pathogens in coastal habitats"

### Supplementary figure 2

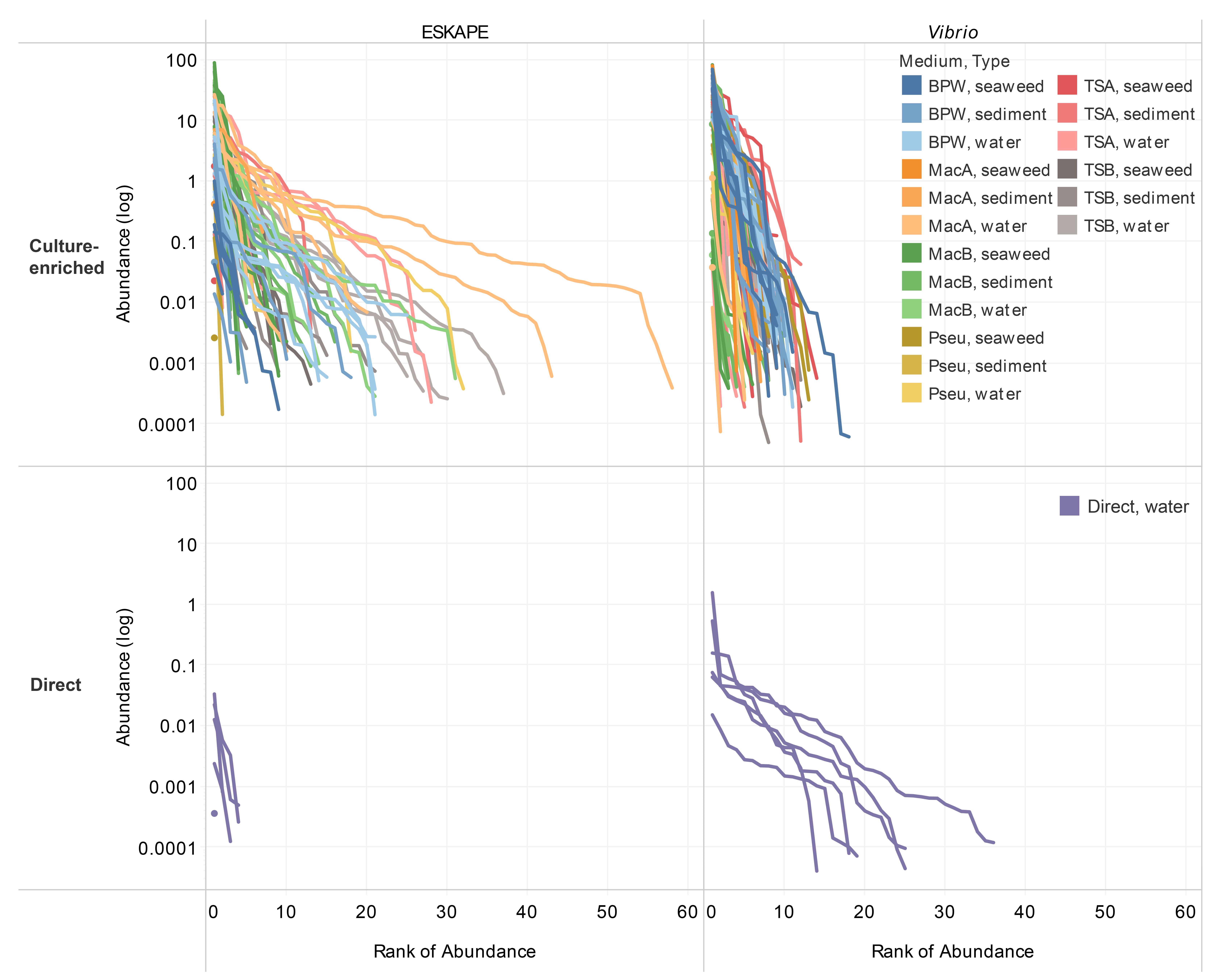

### Supplementary figure 3

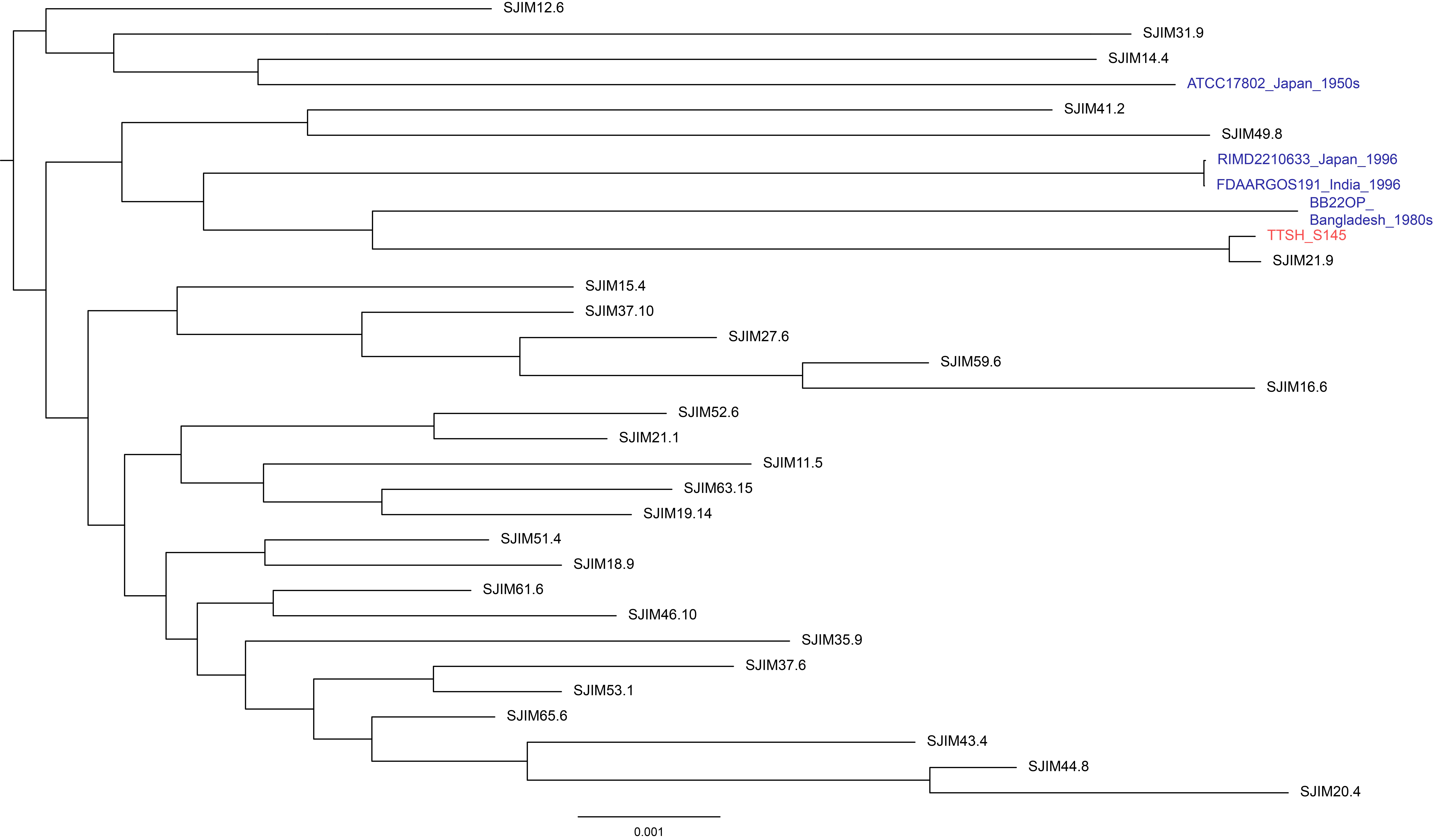
